# Oxygen dependent mitochondrial formate production and the reverse Pasteur effect

**DOI:** 10.1101/2020.04.10.035675

**Authors:** Johannes Meiser, Alexei Vazquez

**Author notes:** Corresponding author, Alexei Vazquez, Cancer Research UK Beatson Institute, Switchback Road, Bearsden, Glasgow G61 1BD, UK.

## Abstract

The Pasteur effect dictates that oxygen induces respiration and represses fermentation. However, we have shown that oxygen stimulates mitochondrial formate production and excess formate production induces glycolysis in mammalian cells. Our observations suggest the hypothesis that increased respiration induces an increase, rather than a decrease, of fermentation, the reverse Pasteur effect. Using a mathematical model we show that, in the absence of mitochondrial formate production, we should always observe the Pasteur effect, a reduction in fermentation with increasing respiration. However, in cells with active mitochondrial formate production, the rate of fermentation first increases with increasing the rate of respiration, indicating a metabolic sweet spot at moderate oxygen availability that is within the range of tissue oxygen tensions. We provide experimental evidence for the manifestation of the reverse Pasteur effect at such oxygen tension.

## Background

Pasteur described fermentation as life without oxygen and postulated its repression by oxygen [1], the Pasteur effect. Yet, Brown observed that yeast cells ferment at a higher rate under the presence rather than absence of oxygen [2]. Later on Warburg reported his observations on aerobic glycolysis by cancer cells [3, 4]. Warburg hypothesized that cancer cells have defective mitochondria and consequently require higher rates of glycolysis to sustain the energy demand of tumour growth [5]. Working with normal tissues subject to viral infections, Crabtree demonstrated that aerobic glycolysis is not unique to cancer cells and he postulated a repression of respiration by glycolysis [6], the Crabtree effect. Today it is well established that aerobic glycolysis is an ubiquitous phenotype of cells from all life kingdoms [7] and cancer cells as well [8, 9].

Aerobic fermentation and the Pasteur effect are not mutually exclusive. Some strains of yeast exhibit aerobic fermentation of glucose to ethanol, the Crabtree positive yeasts. Yet, their fermentation rate increases further when exposed to reduced oxygen tensions [10]. In the context of mammals, it is well known that fermentation of glucose to lactic acid is increased when tissues are subject to hypoxia [11–13]. Looking at it the other way around, an increase in the oxygen tension decreases the rate of fermentation as postulated by the Pasteur effect. What is peculiar about aerobic fermentation is that the repression is not complete. Aerobic fermentation in cells with active oxidative phosphorylation is an incomplete Pasteur effect.

We have recently shown that mitochondrial formate production from the catabolism of serine can also stimulate glycolysis [14]. Since the rate of serine catabolism to formate increases in proportion to the rate of oxidative phosphorylation [15, 16], this new evidence leads to a conundrum. Increased oxidative phosphorylation relieves cells from the necessity of glycolysis to satisfy their energy demand, thus resulting in a decrease of fermentation, or lactate release in mammalian cells. In contrast, increased oxidative phosphorylation induces an increase in the mitochondrial serine catabolism to formate that promotes glycolysis [14].

Here we provide theoretical and experimental evidence that these two opposite tendencies result in an unanticipated phenotype, the increase of fermentation when increasing the availability of oxygen. We call this phenotype the reverse-Pasteur effect.

## Methods

### HAP1 cells

Formate release rates were obtained from a previous report [16]. Previously derived raw LC-MS data for HAP1 cell cultures supernatant [16] was re-analysed with TraceFinder (Thermo Fisher Scientific) software to extract and quantify the lactate peaks. The exchange rate of lactate was then estimated as described previously [15].

### Yeast cells

The data for yeast cells was obtained from Table 1 in reference [10] without any additional processing.

### Statistical analysis

The experimental data for the HAP1 cells was derived from three independent experiments, each with three technical replicates. The statistical significance was calculated with a Welch’s *t*-test, two tails and unequal variance. The experimental data for yeast cells just included mean and standard deviations and no statistical test was performed.

**Figure 1.**
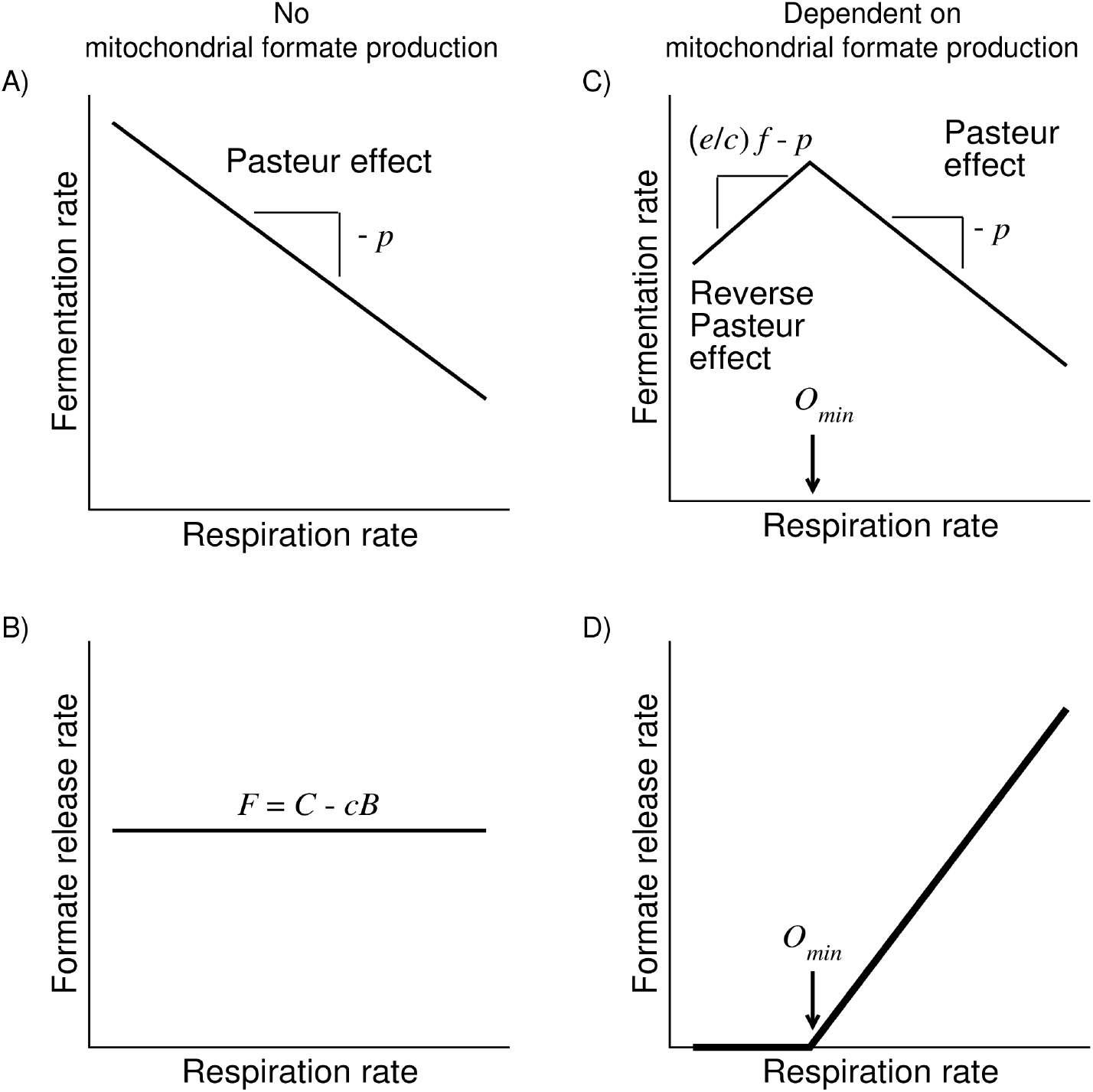
Predicted scenarios. Predicted rates of fermentation and formate release depending on the status of mitochondrial formate production.

## Results

### Mathematical model

We consider a simplified mathematical model accounting for the balance of energy and one-carbon units. (1) The energy balance includes energy production from glycolysis (substrate phosphorylation) and respiration (oxidative phosphorylation) and energy consumption by cell maintenance and biosynthesis. (2) The one-carbon units balance takes into account the one-carbon unit production by cytosolic one-carbon metabolism and mitochondrial formate production, the one-carbon unit demand of biosynthesis and formate release. These balances are mathematically represented by the equations

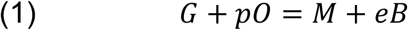

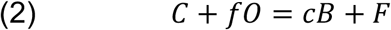

Here *G* is the rate of ATP production by glycolysis coupled to fermentation, *p* is the pO ratio of oxidative phosphorylation, *O* is the rate of respiration, *M* is energy demand for cell maintenance, *e* is the energy demand of biosynthesis and *B* is the biosynthesis rate. *C* is the rate of one-carbon production from the cytosolic pathway, *f* is the mitochondrial formate production rate per unit of oxygen consumed, *c* is the one-carbon unit demand to double the cell content and *F* is the rate of formate release.

### Fermentation rate without mitochondrial formate production

We first consider a scenario where there is no mitochondrial formate production (*f* = 0) and the cytosolic one-carbon metabolism is the only source of one-carbon units. In this case our equations (1) and (2) simplify to

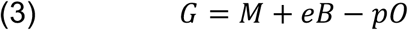

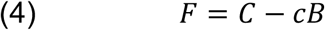

In this case the rate of glycolysis decreases with increasing rate of respiration, recapitulating the Pasteur effect (**Fig. 1A**). In this case, the rate of formate release is predicted to be constant and independent of the respiration rate (**Fig. 1B**). We note *F* can be zero, meaning no formate release, if biosynthesis is limited by the rate of one-carbon unit production from the cytosolic pathway (*C* = *cB*).

### Fermentation rate dependent on mitochondrial formate production

We next consider a scenario where mitochondrial formate production is essential to sustain the one-carbon unit demand of biosynthesis (*C* < *cB*). In this case cells require a minimum rate of respiration (*O_min_*) to satisfy the one-carbon demand of biosynthesis. *O*_min_ is calculated from equation (2) setting the formate release F to zero, resulting in

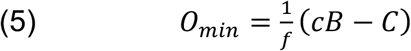

When the respiration rate is above this minimum (0 > *O_min_*) biosynthesis is not limited by the availability of one-carbon units, although still dependent on mitochondrial formate production. In this case the rates of lactate and formate release are independent from each other and can be calculated solving for *G* and *F* in equations (1) and (2), respectively.

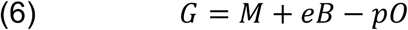

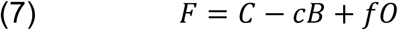

In this case the fermentation rate decreases with increasing the rate of respiration, once again recapitulating the Pasteur effect (**Fig. 1C**, to the right of *O_min_*). In contrast, due to the excess one-carbon unit production, the formate release rate increases with increasing the rate of respiration (**Fig. 1D**, to the right of *O_min_*).

The picture changes dramatically when the respiration rate falls below the minimum (0 < *O_min_*). Here biosynthesis is limited by the availability of one-carbon units, there is no formate release and the fermentation and biosynthesis rates are obtained solving equations (1) and (2) for *G* and B, resulting in

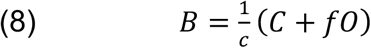

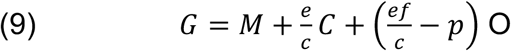

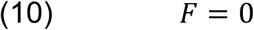

The inspection of equation (9) suggests two different scenarios. When *ef < pc* the fermentation rate will again decrease with increasing the respiration rate. In contrast, when *ef* > *pc*, we uncover the unanticipated possibility that the fermentation rate increases with increasing the respiration rate (**Fig. 1C**, left of *O*_min_). We term this scenario the reverse Pasteur effect, given that the lactate release increases rather than decreases with increasing respiration. This phenomenon is predicted to happen when cells transition from close to anoxic to moderate hypoxic conditions (**Fig. 1C**, left of *O*_min_). When oxygen tension increases further and exceeds the predicted threshold (*O*_min_), then the Pasteur effect occurs. We also note that in the context of the reverse Pasteur effect the rate of biosynthesis increases in proportion to the rate of respiration, as dictated by equation (8).

**Figure 2.**
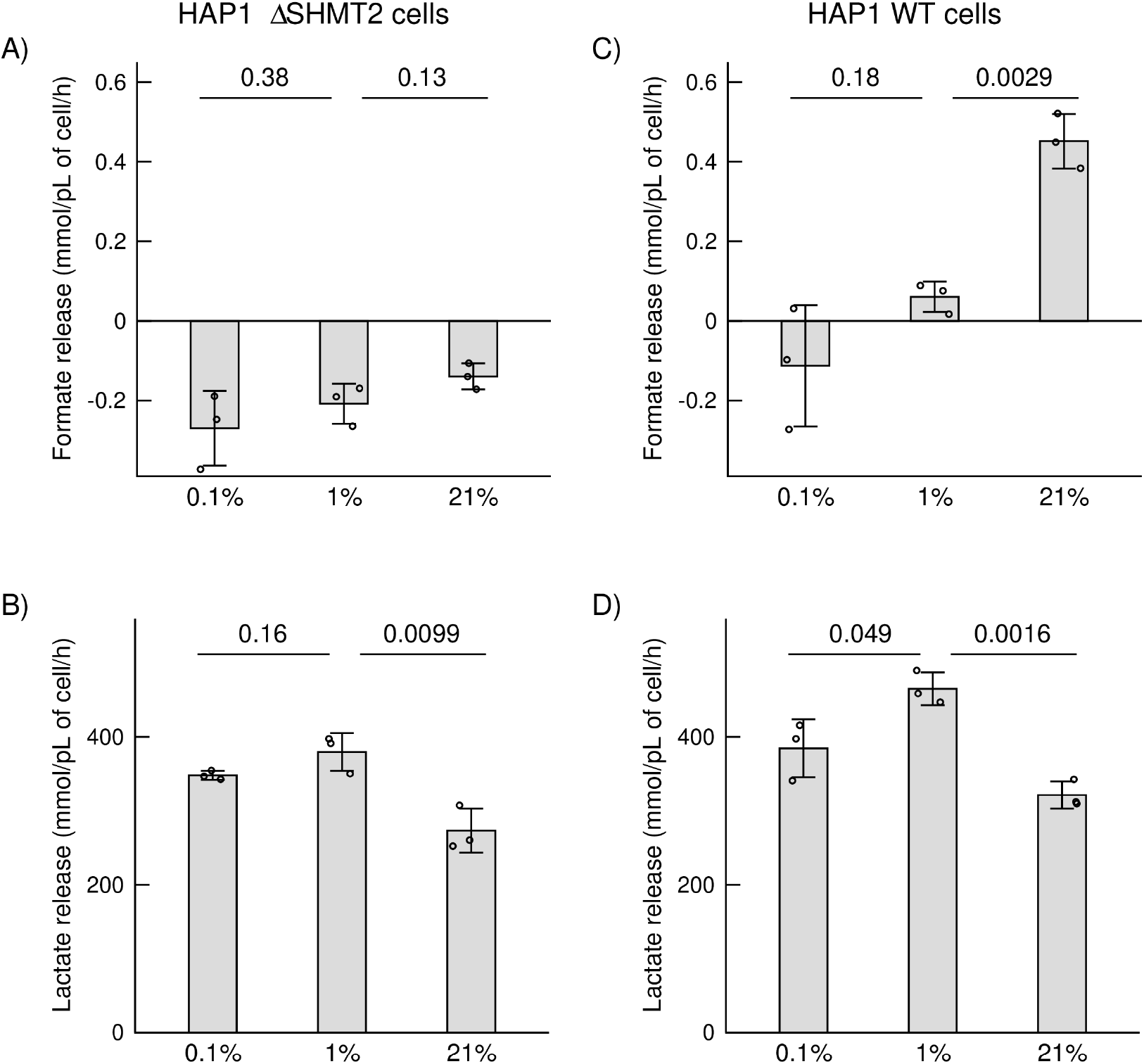
Reverse Pasteur effect in HAP1 cancerous cells. Formate and lactate release rates for ΔSHMT2 and WT HAP1 cancerous cells. Bar heights represent the mean, error bars the standard deviation and symbol independent experiments (triplicate wells per experiment). The figures over the line indicate the statistical significance of differences between groups, Welch’s *t*-test, two tails and unequal variance.

### Experimental verification

To test the theoretical predictions, we have analysed previously reported data for wild-type HAP1 cells (WT) and HAP1 cells with genetic inactivation of mitochondrial serine hydroxymethyl transferase (ΔSHMT2), cultured WT and ΔSHMT2 cells under normoxia, moderate hypoxia and severe hypoxia (21, 1 and 0.1% oxygen) [16]. The formate release rates were extracted directly from the previous report. The lactate release rates are reported here for the first time, after re-analyzing the raw LC-MS data quantifying media metabolites.

ΔSHMT2 cells do not have mitochondrial serine catabolism to formate, representing the case where the cytosolic pathway satisfies the one-carbon unit demand of cell growth (**Fig. 1 A, B**). ΔSHMT2 cells do not release formate (**Fig. 2A**), suggesting that these cells are limited by the availability of one-carbon units (*C* = *cB* scenario). Based on this evidence, our model predicts that the fermentation rate by ΔSHMT2 cells should decrease with increasing the respiration rate. This is indeed what we observe experimentally. At 0.1 and 1% oxygen, ΔSHMT2 cells have similar rates of lactate release, the product of mammalian cell fermentation. But their rate of lactate release drops significantly at 21% oxygen (**Fig. 2B**).

**Figure 3.**
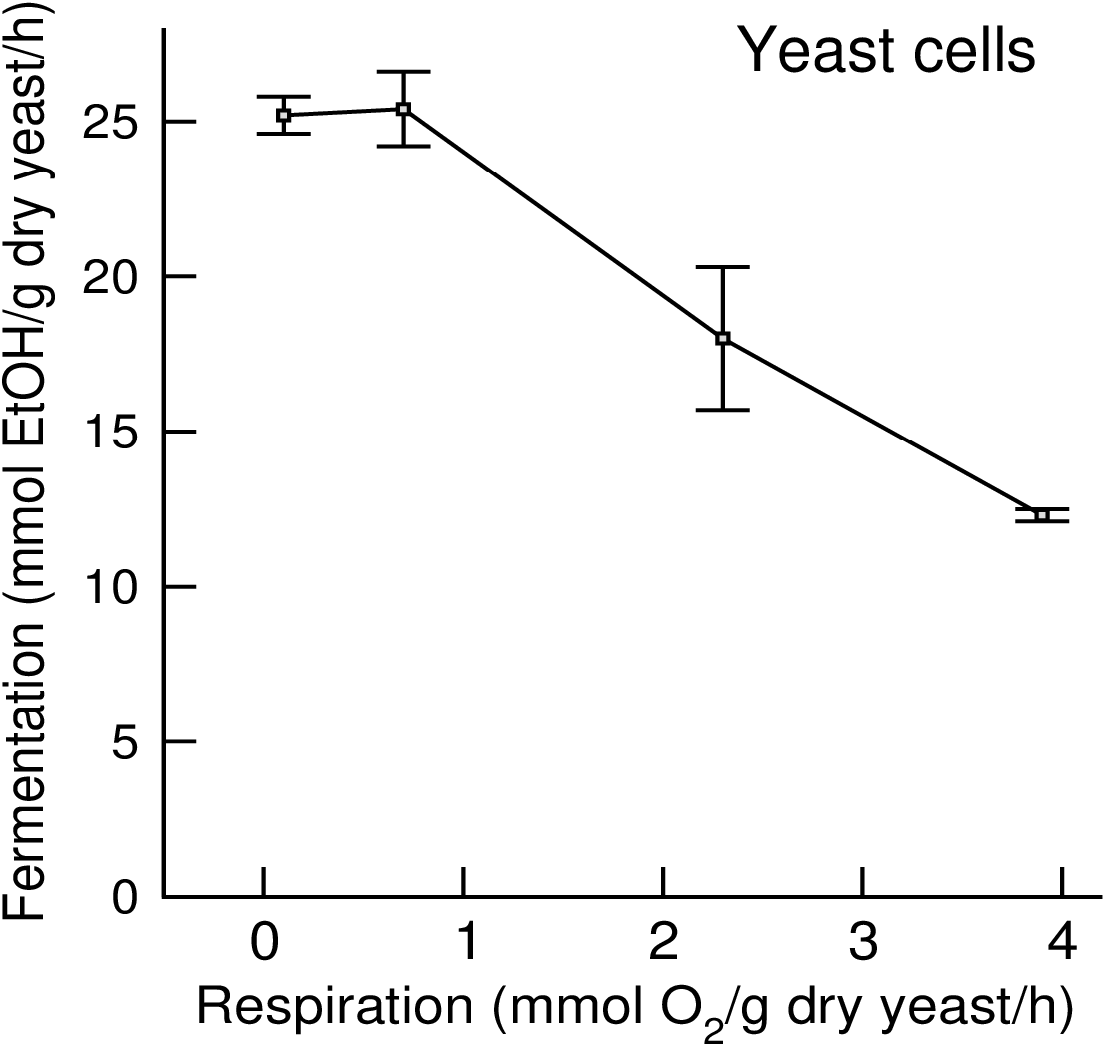
Reverse Pasteur effect in yeast. Rate of fermentation by yeast cells as a function of the rate of respiration. Symbols represent the mean and error bars the standard deviation.

WT cells exhibit a formate release pattern that matches exactly what is expected in cells dependent on mitochondrial formate production (**Fig. 1D**). WT cells do not release formate at 0.1%, a small amount at 1% and they have a significant formate release rate at 21% oxygen (**Fig. 2B**). These cells are candidates to manifest the reverse Pasteur effect. We observe that this is indeed the case. WT cells significantly increase their rate of lactate release from 0.1% to 1% oxygen, but significantly decrease their rate of lactate release from 1% to 21% oxygen. This data demonstrates that, in cells with active mitochondrial formate production, the rate of lactate release is maximum at an intermediate oxygen level.

The compartmentalization of one-carbon metabolism into mitochondrial formate production and cytosolic one-carbon metabolism is conserved from mammalian cells to yeast cells [17]. Yeast cells can also exhibit formate overflow [18], indicating that they can be in a metabolic state of excessive one-carbon unit production. Yeast is therefore another candidate system to manifest the reverse Pasteur effect. We have collected previously reported data for yeast cells using glucose as their carbon source [10]. That included the rate of ethanol release, the fermentation product of yeasts, and the rate of respiration. At high respiration rates there is an evident Pasteur effect with a characteristic decrease in the ethanol release rate with increasing rate of respiration (**Fig. 3**). However, the data for the two lowest respiration rates deviates from this behavior. For these two points the ethanol release rate is approximately constant with increasing the respiration rate. This behavior is what the model predicts for a case where the tendency to increase fermentation due to increased mitochondrial formate production balances the increase in energy generation from oxidative phosphorylation (*ef* = *cp* case of the mathematical model).

## Discussion

We have provided theoretical support for a new phenotype, the reverse Pasteur effect, whereby the rate of fermentation increases rather than decreases with increasing rate of respiration. This effect manifests at low oxygen tensions when cells transition from nearly anoxic to mild hypoxic oxygen concentrations. When oxygen concentration increases further, the classical Pasteur effect can be observed. To validate this theoretical prediction, we provide two experimental systems manifesting this effect: the mammalian cancer cell line HAP1 and yeast cells. We predict that the reverse Pasteur effect is a general feature of eukaryote cells with mitochondrial formate production coupled to respiration.

In the context of mammalian physiology, the reverse Pasteur effect forces us to reconsider the relation between lactate accumulation and tissue oxygenation. The oxygen tension of mammalian tissues is in the range of 1-5% under normal physiological conditions. That is the range where the HAP1 cells manifest their highest rate of lactate production. Physiological hypoxia would be a reduction in the oxygen tension down to values around 0.1%. In this context, the decision whether cells increase or decrease their rate of fermentation would depend on whether they have a biosynthetic demand for one-carbon units and whether they are dependent on mitochondrial formate production to satisfy that demand. In the absence of a biosynthetic demand for one-carbon units, lactate production should increase when going from 1% to 0.1% oxygen, as generally expected. When there is a biosynthetic demand of one-carbon units that is dependent on mitochondrial formate production, the reverse Pasteur effect should manifest as a decrease in lactate production when going from 1% to 0.1% oxygen.

The mitochondrial formate production is essential for the closure of the neural tube during embryonic development [19]. This provides a context where the reverse Pasteur effect could play a physiological role. During embryonic development there is a complex interplay between cell proliferation and the development of the vascular system, the latter providing the oxygen required for cell proliferation. Whether the reverse Pasteur effect constitutes a mechanism with *in vivo* relevance to couple oxygen availability with proliferation remains to be determined.

We anticipate the relevance of the reverse Warburg effect in the context of ischemia and reperfusion [20]. Our prediction would be that glycolysis increases from ischemia to the initial stages of reperfusion. Determining whether that is the case is complicated by the fact that circulating lactate is a substrate of oxidation phosphorylation [21]. Therefore, changes in the circulating lactate concentration could be due to changes in lactate production by glycolysis or changes in lactate oxidative phosphorylation and both are modulated by oxygen availability.

Our analysis expands the richness of mixed respiratory-fermentation phenotypes. The Pasteur effect dictates that respiration represses fermentation. The Warburg effect tells us that this repression may not be complete and cells may exhibit fermentation even when there is plenty of oxygen available. The reverse Pasteur effect provides an additional correction when oxygen tensions are low, pointing out that respiration increases fermentation when oxygen availability is increased from very low to intermediate.

## Acknowledgements

This work was supported by Cancer Research UK C596/A21140. JM was supported by a DFG Fellowship (Grant Number ME 4636/2-1) and by a FNR ATTRACT fellowship (Grant Number: A18/BM/11809970).

